# Oxidative stress biomarkers in hypothyroid, non thyroid illness and euthyroid dogs

**DOI:** 10.1101/2023.08.02.551618

**Authors:** K Mohanambal, K Satish Kumar, P Nagaraj, M Usha Rani, Y Ravi Kumar

## Abstract

There are only a few numbers of published reports available on oxidative stress parameters in hypothyroidism and many of which are in the field of human medicine. Studies on hypothyroidism in dogs lack an assessment of oxidative stress and some are vague and inconclusive. So, the current study was formulated primarily to address variations in chief antioxidant parameters in dogs with hypothyroidism, non-thyroid illness and euthyroidism. Secondly, magnitude of oxidative stress was measured in hypothyroid dogs prior to and during the course of levothyroxine therapy, which were compared statistically. Evaluation of thyroid profile (TT4, fT4 and cTSH) was carried out prior to the assessment of oxidative stress biomarkers such as, serum malondialdehyde (MDA), superoxide dismutase (SOD), reduced glutathione (GSH) and catalase. A total of 45 dogs were employed in the study, that includes 19 dogs with hypothyroidism, 11 dogs with non-thyroid illness and 15 dogs with euthyroidism. These dogs were brought either to the dermatological unit or the immunization unit. Dogs were divided into three groups: group 1 represented hypothyroidism (low fT4 or low TT4 and high cTSH); group 2 represented non-thyroid disease (low or normal TT4 and low cTSH i.e., inconclusive levels of thyroid hormones); and group 3 represented dogs with normal thyroid levels (euthyroid dogs). Descriptive statistics and normality plots were carried out using SPSS software version 21. Assuming that the variances were unequal, P values were calculated and compared by one-way ANOVA, post hoc multiple comparison analysis using the Games-Howell test. The mean thyroid hormone levels of hypothyroid dogs before (Day 0) and after (Day 60) treatment was significantly different (P<0.05). Hypothyroid dogs (group 1) displayed significantly lower mean fT4 levels (0.56±0.07) and higher mean cTSH levels (8.14±1.17) when compared to other groups. The mean values of serum biochemical parameters showed significant differences among groups (P<0.01). Various biomarkers showed a significant alteration viz., there was a significant reduction in catalase, SOD, GSH and increase in MDA in hypothyroid dogs when compared with euthyroid dogs. However, following therapy for 60 days, catalase, SOD and GSH levels elevated and MDA level reduced significantly and reached nearly to that of euthyroid dogs. Further, there was no significant difference between NTI and euthyroid dogs which demonstrated the antioxidant defence against oxidative stress in hypothyroid dogs. As an outcome, antioxidant measures and serum biochemical tests can be employed as a diagnostic tool to rule out oxidative stress in hypothyroid dogs.

## Introduction

Oxidative stress arises due to over production of pre-oxidants, sequential damage to the antioxidant system which leads to the formation of free radicals and consecutively accelerates the accumulation of reactive oxygen species (ROS) inside the cells [1]. Intense and prolonged stress gives rise to serious cell damage. Hypothyroidism is one of the most common endocrine dermatoses [2], presented as primary hypothyroidism associated due to lymphocytic thyroiditis or idiopathic thyroid atrophy resulting in degeneration of thyroid gland [3,4]. Oxidative stress parameters (detecting protein damage, lipid damage and DNA damage) were detected directly by enzymatic antioxidant detection and non-enzymatic antioxidant detection methods [5]. Thyroid hormones had close association with oxidative stress [1]. Thyroid hormone is playing an important role in metabolism and in hypothyroidism, increased lipid peroxidation [6] and low antioxidants levels were documented [7]. A reduction in superoxide dismutase activity (SOD), catalase activity and an overall increase in thiobarbituric acid reactive substances (TBARS) in hypothyroid patients were observed [8]. Few authors [9,10] suggested the usage of oxidative biomarkers (malondialdehyde, total antioxidant capacity, TBARS) as a diagnostic tool for hypothyroidism in dogs. Levothyroxine treatment attenuated the changes occurred due to oxidative stress [8]. Most of the time serum and less frequently, saliva was used to study and evaluate the ooxidative status of patients [11–14]. Only few studies related to detect oxidative stress parameters in hypothyroid dogs were available and were inconclusive. Thus, the goal of the current study is to evaluate oxidative biomarkers such as malondialdehyde (MDA), superoxide dismutase (SOD), catalase activity and glutathione reductase (GSH) in dogs with hypothyroidism, non-thyroid illness and euthyroidism. The study also assessed the alterations of antioxidant profile among hypothyroid dogs before and after levothyroxine administration.

## Materials and methods

### Selection of dogs

Forty-five adult treatment naive dogs of different age, breed and gender presented to the small animal dermatology unit and immunization unit of the College of Veterinary Science, Hyderabad were considered for the study. Out of these, 30 were showing classical signs of hypothyroidism and whereas 15 dogs were apparently healthy. All these dogs including apparently healthy dogs were subjected for thyroid profile viz., fT4, TT4 and cTSH to confirm hypothyroidism following standard procedure [10]. Further, dogs with concurrent illness like diabetes mellitus, renal impairment, hyperadrenocorticism and heart failure were excluded from the study. Euthyroid dogs (n=15) were kept as control that were selected based on the following criteria, 1. vaccinated dog above four-year age, 2. healthy individual as per history and clinical examination and 3. normal levels of thyroid profile on assessment. Selected dogs were designated into three groups namely group 1 (hypothyroid dogs), group 2 (NTI) and group 3 (euthyroid).

### Assessment of body condition score (BCS)

The assessment of body condition score (BCS) was done as per the standard procedure [16] with a BCS 9 score scale (1-3 thin, 4 and 5 ideal, 6-9 overweight or obese).

### Laboratory analysis

The whole blood was collected from the patient in the morning hours in a clot activator coated sterile serum vials and kept undisturbed for 15-30 minutes. Further, the vials were centrifuged (1,000-2,000 x g for 10 minutes) for serum separation. Then the serum was aliquoted into labelled eppendorf tubes and stored in deep freezer (-20°C) for the assessment of serum biochemical parameters, thyroid profile and oxidative parameters. The biochemical parameters such as total cholesterol, triglycerides, high density lipoprotein cholesterol (HDL), low density lipoprotein cholesterol (LDL), alanine aminotransferase (ALT), aspartate aminotransferase (AST) and alkaline phosphatase (ALP) were analysed using EM DENSITY 180 fully auto biochemical analyser. Serum thyroid profile such as total thyroxine (TT4), free T4 (fT4) and canine specific thyroid stimulating hormone (cTSH) were analysed [17] and the results were expressed as ng/dL, pg/ml and ng/ml, respectively by using ELISA kit (Fine test, Wuhan). Oxidative parameters measured were, estimation of end product of lipid peroxidation, i.e., malondialdehyde and enzymatic antioxidants such as activity of catalase, superoxide dismutase (SOD) and reduced glutathione (GSH).

#### Estimation of serum thyroid profile

Serum levels of canine TT4 (Thyroxine), fT4 and cTSH (canine specific TSH) were measured by using Fine test ELISA kit with diagnostic sensitivity of 0.422 ng/ml, 0.938 pg/ml and 0.75 ng/ml, respectively. Detection limits were 0.703 - 45 ng/ml, 1.563 - 100 pg/ml and 1.25 - 80 ng/ml, respectively for the above tests. Intra and inter assay covariance for all the assays were <8% and <10%. No significant cross reactions between thyroid hormones and analogues were observed. The standard curve was generated as per the kit standards [17].

#### Estimation of Catalase activity

The Catalase activity of a given sample was measured by calculating the amount of hydrogen peroxide micro molecules consumed per minute per micro gram of sample [18]. A 2 ml eppendorf tube was filled with 1 ml of 0.1 M PBS, followed by 1 ml EDTA and 50 µl of RBC sample. All the tubes were incubated at room temperature for 10 minutes and later added with 950 µl H_2_O_2_ solution. The rate of change of absorbance per minute was measured at 240 nm. Micro moles of H_2_O_2_ consumed/min/mg was calculated by OD of sample x 1000 divided by 43.6 x 1.5 (43.6 and 1.5 are extinction coefficients). Units of activity was represented as U/mg protein.

#### Estimation of superoxide dismutase (SOD)

The serum activity of SOD was measured by microtiter plate assay method [19]. In this method, the production of superoxide by pyrogallol auto oxidation was done by tetrazolium dye [MTT (3-(4,5-dimethyl-thiazol-2-yl) 2,5-diphenyl tetrazolium bromide)]. The above reaction converts the colourless substrate into a coloured (yellow) product. Dimethyl sulfoxide (DMSO) was added to terminate the reaction and colour produced remained stable. Increased level of SOD in the samples lead to the reduction of superoxide concentration. In each well of a flat-bottomed microtiter plate, 6 µl of MTT was added, followed by 100 µl of serum sample (in duplicate) and 15 µl of pyrogallol. The positive control was 15 µl pyrogallol and 29 µl of PBS (Phosphate buffered saline) and negative control was 44 µl of PBS. The plate was incubated at room temperature for 10 minutes then 150 µl of DMSO was added to all the wells to stop the reaction. The plate was read by ELISA reader at 570 nm wavelength. The SOD was calculated by the following formula, SOD = positive control OD - Negative control OD x 17 x 0.5. Unit of activity was represented by U/mg of protein.

#### Estimation of Glutathione (GSH)

Glutathione reductase catalyses the reductions of oxidized glutathione producing a reduced glutathione by consuming NADPH or NADH. The reaction was irreversible. Enzyme activity was estimated by measuring the rate of decrease in OD resulted by the oxidation of NADPH [20]. The cuvette was filled with 2 ml of 0.1M PBS buffer followed by 100 µl of serum. Later, 100 µl of glutathione oxidised solution was further added with 100 µl of FAD solution and 100 µl of EDTA, respectively. Subsequently, incubation was done for 15 minutes at room temperature. Later, 100 µl of NADPH solution was added. Finally, 300 µl of supernatant was collected from the cuvette and added to 96 well flat bottom ELISA microtiter plate. The OD was read at 340 nm wavelength. The GSH was calculated by the following formula, GSH = OD x 3781 (3781 was the extinction coefficient). Unit of activity was expressed as U/ml.

#### Estimation of malondialdehyde

Malondialdehyde (MDA), is a secondary product of lipid peroxidation as well as useful marker to evaluate the extent of lipid peroxidation in the sample. MDA reacts with 2-thio barbituric acid and forms trimethyne, a pink coloured substance, that absorbs light at 535 nm [21]. In a clean test tube 100 µL of distilled water and 50 µl of sodium dodecyl sulphate (SDS 8% W/V) was added and kept as blank. In subsequent test tubes, 100 µL of serum, 100 µL of distilled water, 50 µl of SDS were added and incubated at room temperature for 10 minutes. Then 375 µl of 20 % acetic acid and 375 µl of TBA were added and placed in boiling water bath for 60 minutes. Further, 2.5 ml of distilled water and 1.25 ml of 15:1 butanol pyridine solution were added and centrifuged for 10 minutes at 3000 rpm. Later, 300 µl of supernatent was collected and added to 96 well flat bottom ELISA microtiter plate. Calculation was done by the formula, total volume in test tube (4.75) x OD of sample/0.156. The results were expressed in nmoles of MDA.

### Ethical statement

This study was approved by animal research ethics committee. The protocol was approved by Institutional Animal Ethics Committee (IAEC), P. V. Narsimha Rao Telangana Veterinary University, Hyderabad, India (Proposal No. 27/26/CVSc, Hyd. IAEC). No surgical procedures were involved.

## Results

### Study parameters

Basic details of the dogs selected for the study are presented in Table 1. The mean age of hypothyroid dogs was 4.92 years. The mean BCS (Fig 1) of hypothyroid dogs (6.2±0.34) were significantly higher than (P<0.05) euthyroid dogs (4.8±0.29).

**Fig 1.**
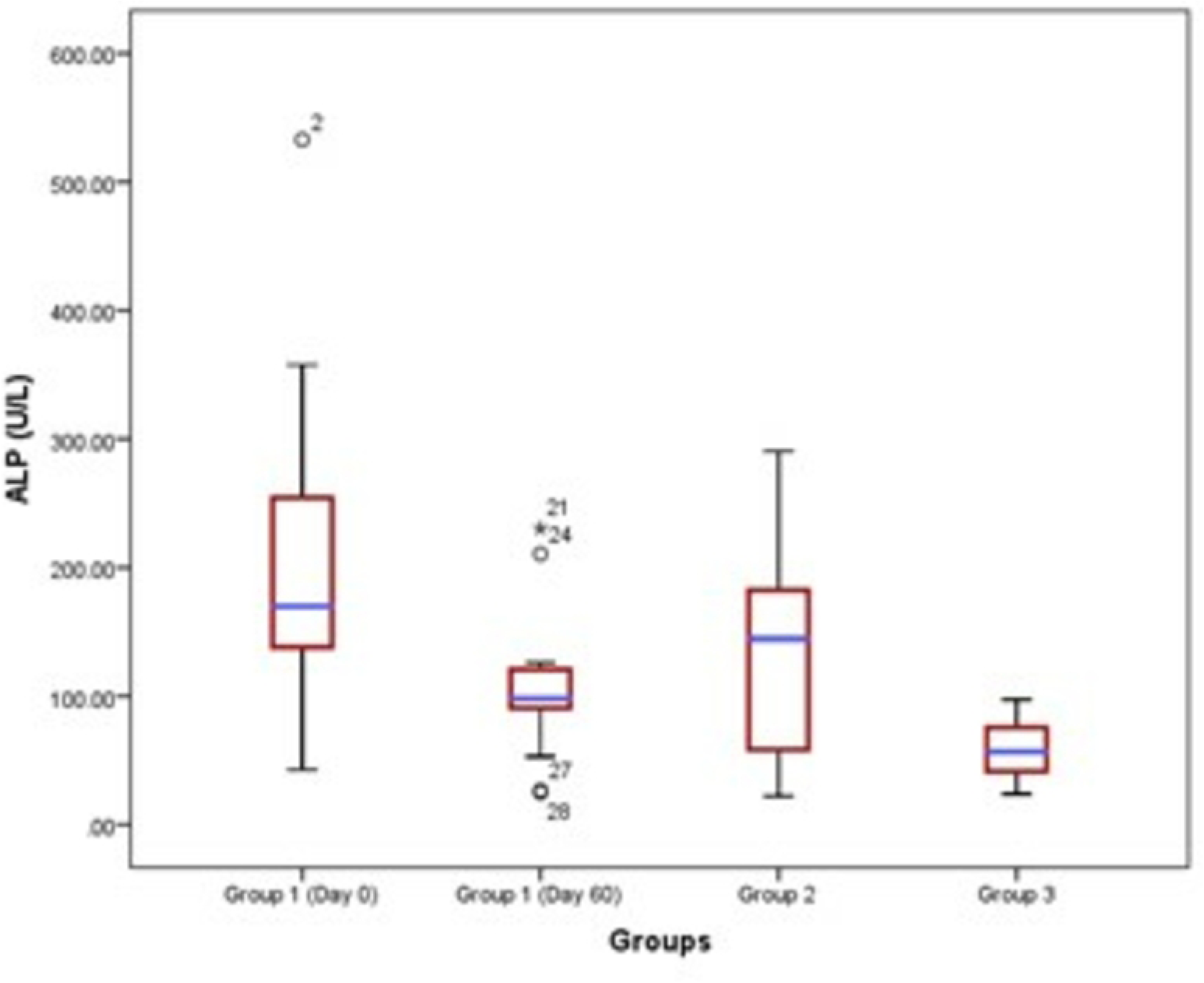
Mean BCS of Hypothyroid, Non thyroid illness and Euthyroid dogs.

**Table 1.** Demographic characteristics of the dogs selected for the study.

| Basic details | Hypothyroid dogs (Group 1) | Dogs with NTI (Group 2) | Euthyroid dogs (Group 3) | F value | Sig. |
| --- | --- | --- | --- | --- | --- |
| N | 19 | 11 | 15 |  |  |
| **Breed** | Beagle-1<br>Chow Chow-1<br>Dachshund-2<br>German Shepherd-2<br>Golden Retriever-2<br>Great Dane-1<br>Labrador (black)-1<br>Labrador (Fawn)-4<br>Lhasa Apso-1<br>Mongrel-1<br>Pomeranian-2<br>Pug-1 | Beagle-1<br>Dachshund-1<br>German Shepherd-1<br>Golden Retriever-2<br>Huski-1<br>Labrador (Fawn)-2<br>Mongrel-1<br>Pomeranian-1<br>Pug-1 | Beagle-1<br>Chow Chow-1<br>Dachshund-1<br>German Shepherd-2<br>Golden Retriever-2<br>Great Dane-1<br>Labrador (black)-1<br>Labrador (Fawn)-2<br>Lhasa Apso-1<br>Mongrel-1<br>Pomeranian-1<br>Pug-1 |  |  |
| **Age**<br>**(Mean±SE)** | 4.92±0.62 | 5.59±1.02 | 5.73±0.88 | 0.317 | 0.730 |
| Minimum age | 1 | 1 | 1.4 |  |  |
| Maximum age | 11 | 13 | 11 |  |  |
| Median age | 4.5 | 6.0 | 6.0 |  |  |
| **Sex** |  |  |  |  |  |
| Male | 11 | 8 | 9 |  |  |
| Female | 8 | 3 | 6 |  |  |
| Male: Female | 1.37:1 | 2.33:1 | 1.5:1 |  |  |
| **BCS**<br>**(Mean±SE)** | 6.2*±0.34 | 5.6±0.27 | 4.8*±0.29 | 5.148 | 0.010 |
| Minimum BCS | 4.0 | 4.0 | 3.0 |  |  |
| Maximum BCS | 9.0 | 7.0 | 7.0 |  |  |
| Median BCS | 6.0 | 6.0 | 5.0 |  |  |

*The mean bearing\*differs significantly at 0.05 level (P<0.05)*

### Evaluation of serum

#### Assessment of serum thyroid profile

The mean fT4, TT4 and cTSH levels of hypothyroid, nonthyroid illness and euthyroid dogs are presented in the Table 2 and Figs (2–4). There is a significant difference (P<0.05) in the mean thyroid hormone levels of hypothyroid dogs before (day 0) and after (day 60) treatment. There is a significant difference (P<0.05) in all the mean thyroid hormone levels of group 1 hypothyroid dogs on day 0 when compared to euthyroid dogs. On day 0 they had a significantly (P<0.05) lower fT4 levels (0.56±0.07) and higher mean cTSH levels (8.14±1.17) compared to other groups. At the same time TT4 levels of group 1 showed highly significant (P<0.05) variation from group 3 but not with group 2. However, the thyroid profile of NTI showed significant difference (P<0.05) in TT4 level with euthyroid dogs. Further, cTSH levels of NTI dogs were non-significant with euthyroid dogs but significant with hypothyroid dogs. At the same time, after levothyroxine treatment (day 60) mean thyroid hormone levels of hypothyroid dogs showed non-significant difference with NTI and euthyroid dogs.

**Fig 2.**
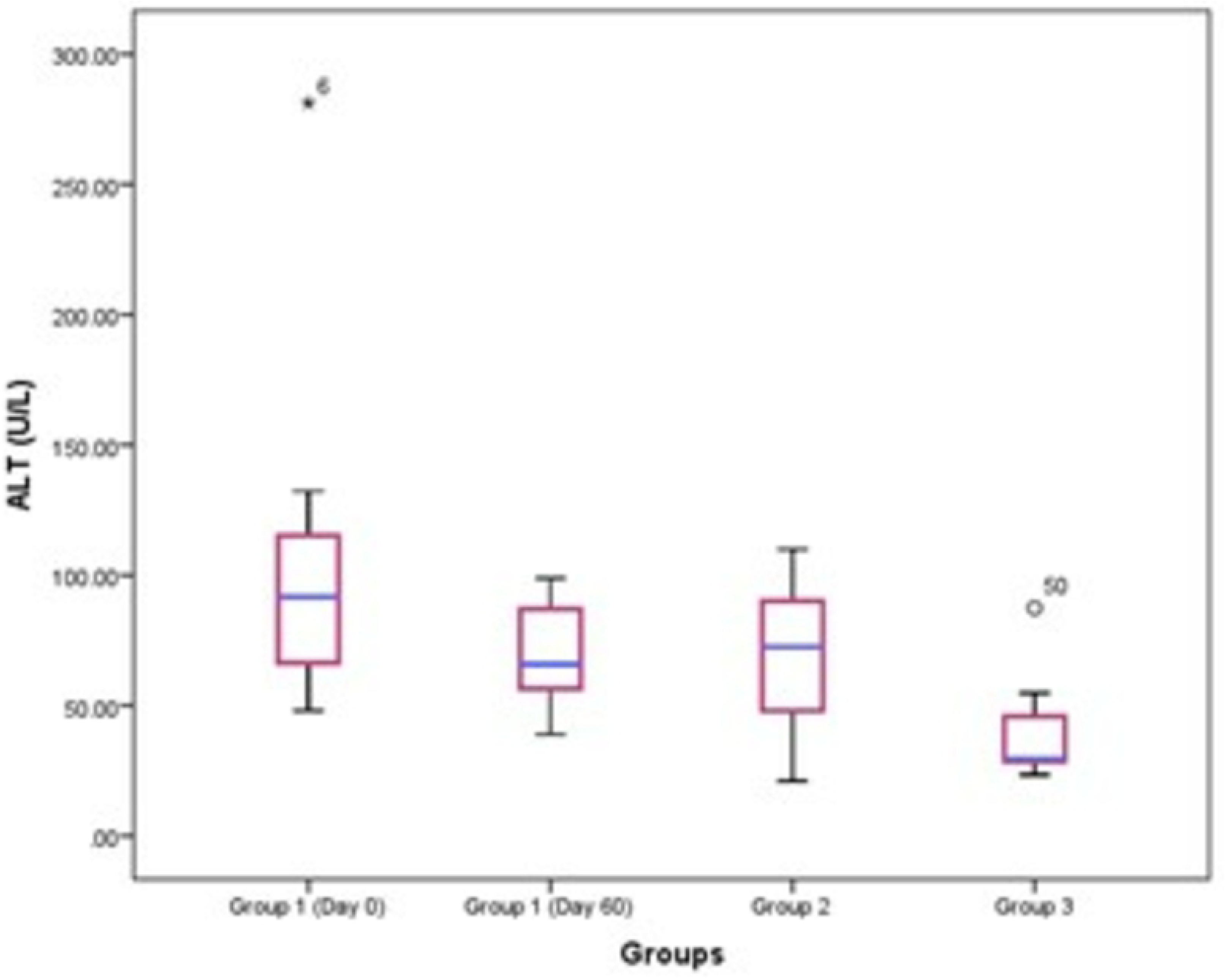
Box plot showing concentration and mean levels of various thyroid hormones in dogs. Concentration and mean levels of serum fT4 in dogs. The box plots showing minimum value, the first quartile, the median, the third quartile and the maximum value (vertical number line of Y axis). A value that lies in both extremes of data marked as dots (outlier values).

**Fig 3.**
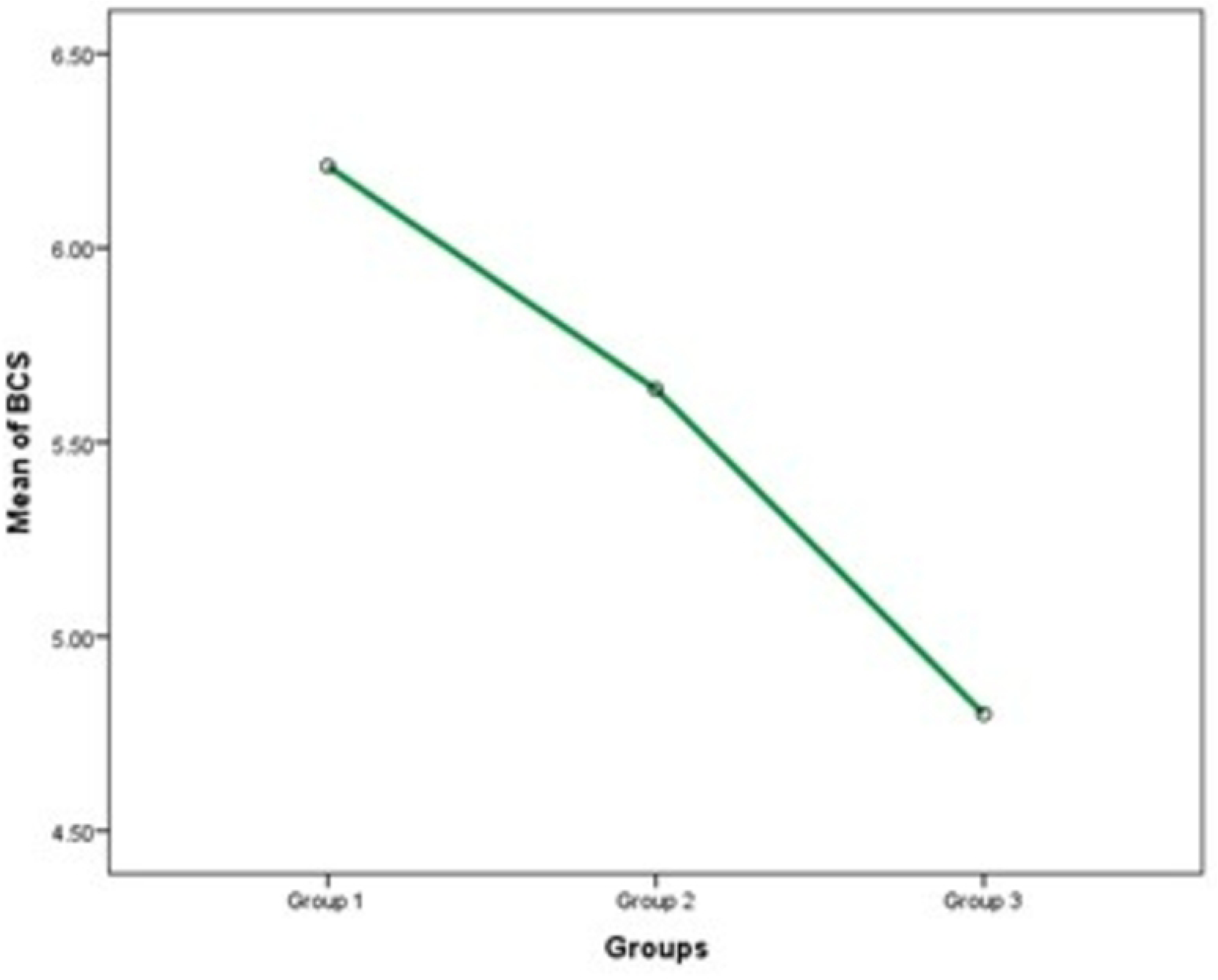
Box plot showing concentration and mean levels of various thyroid hormones in dogs. Concentration and mean levels of serum TT4 in dogs. The box plots showing minimum value, the first quartile, the median, the third quartile and the maximum value (vertical number line of Y axis). A value that lies in both extremes of data marked as dots (outlier values).

**Fig 4.**
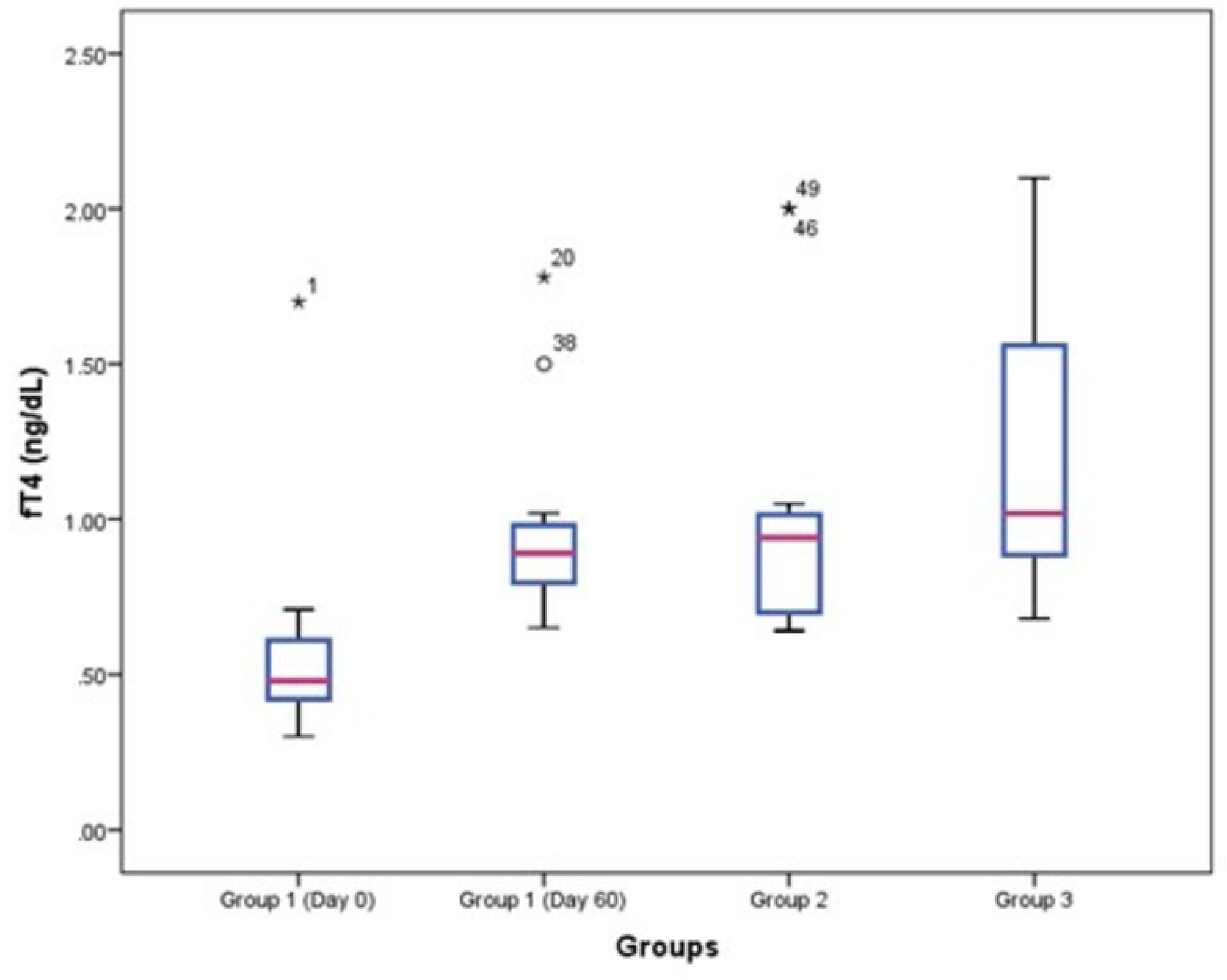
Box plot showing concentration and mean levels of various thyroid hormones in dogs. Concentration and mean levels of serum cTSH in dogs. The box plots showing minimum value, the first quartile, the median, the third quartile and the maximum value (vertical number line of Y axis). A value that lies in both extremes of data marked as dots (outlier values).

**Table 2.**
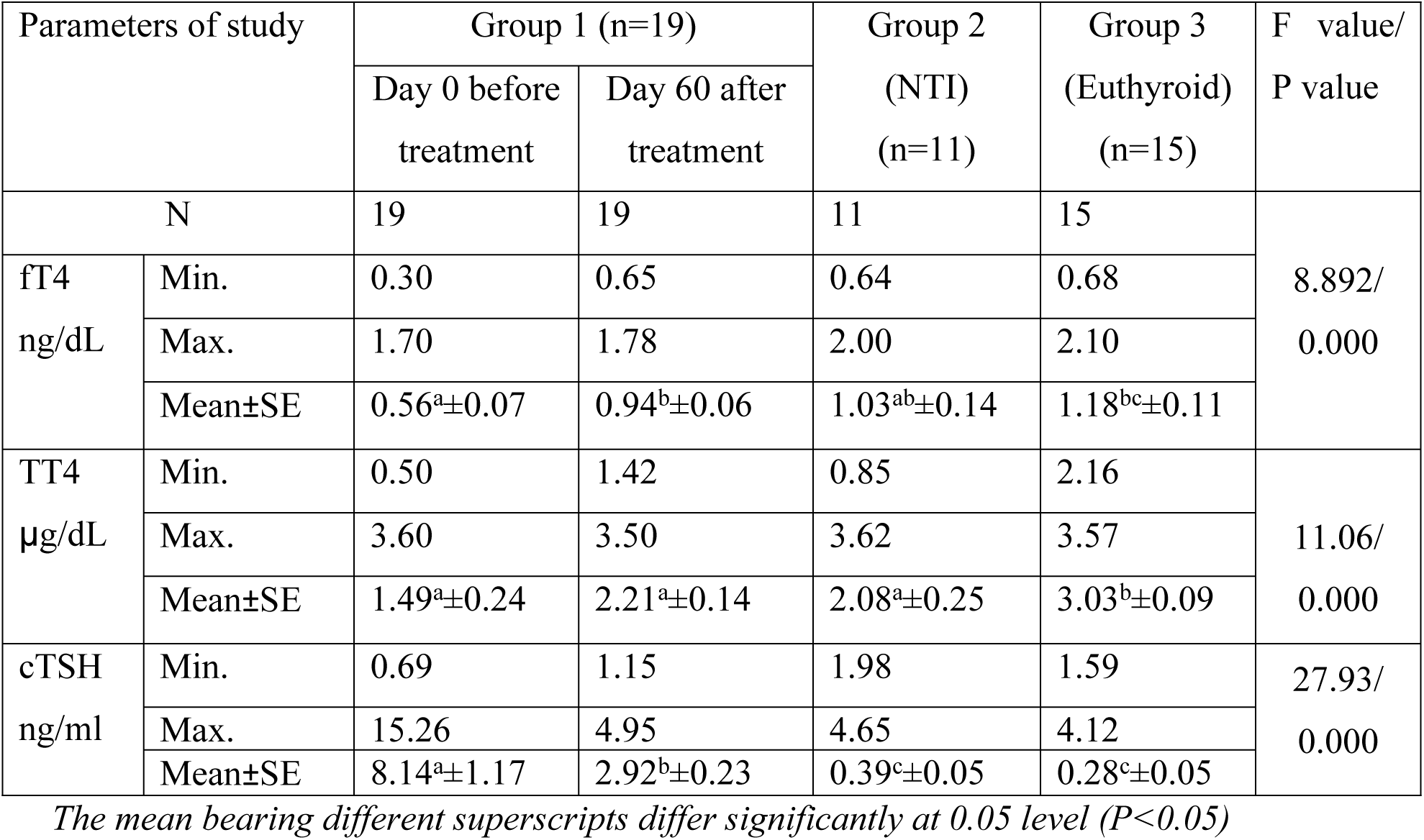
Mean serum thyroid profile of dogs of various groups selected for the study.

#### Assessment of serum biochemical parameters

Serum biochemical parameters such as ALP, ALT, AST, cholesterol, triglyceride, HDL and LDL were analyzed in hypothyroid, non thyroid illness and euthyroid dogs (Table 3). ALP, ALT, AST, cholesterol, triglycerides, HDL and LDL levels of group 1 (day 0) dogs showed highly significant variations in mean values compared to group 3 (P<0.05). Further, AST, HDL and LDL levels of group 1 (day 0) showed significant changes in mean values compared to group 2 (P<0.05). Further, within group 1 dogs, levels of ALP, AST, cholesterol, HDL and LDL varied significantly before (day 0) and after (day 60) therapy (P<0.05). Invariably, the mean values of all the biochemical parameters studied in hypothyroid dogs showed a highly significant changes following treatment for 60 days, which outlined the importance of levothyroxine therapy in canine hypothyroidism (Figs 5–11).

**Fig 5.**
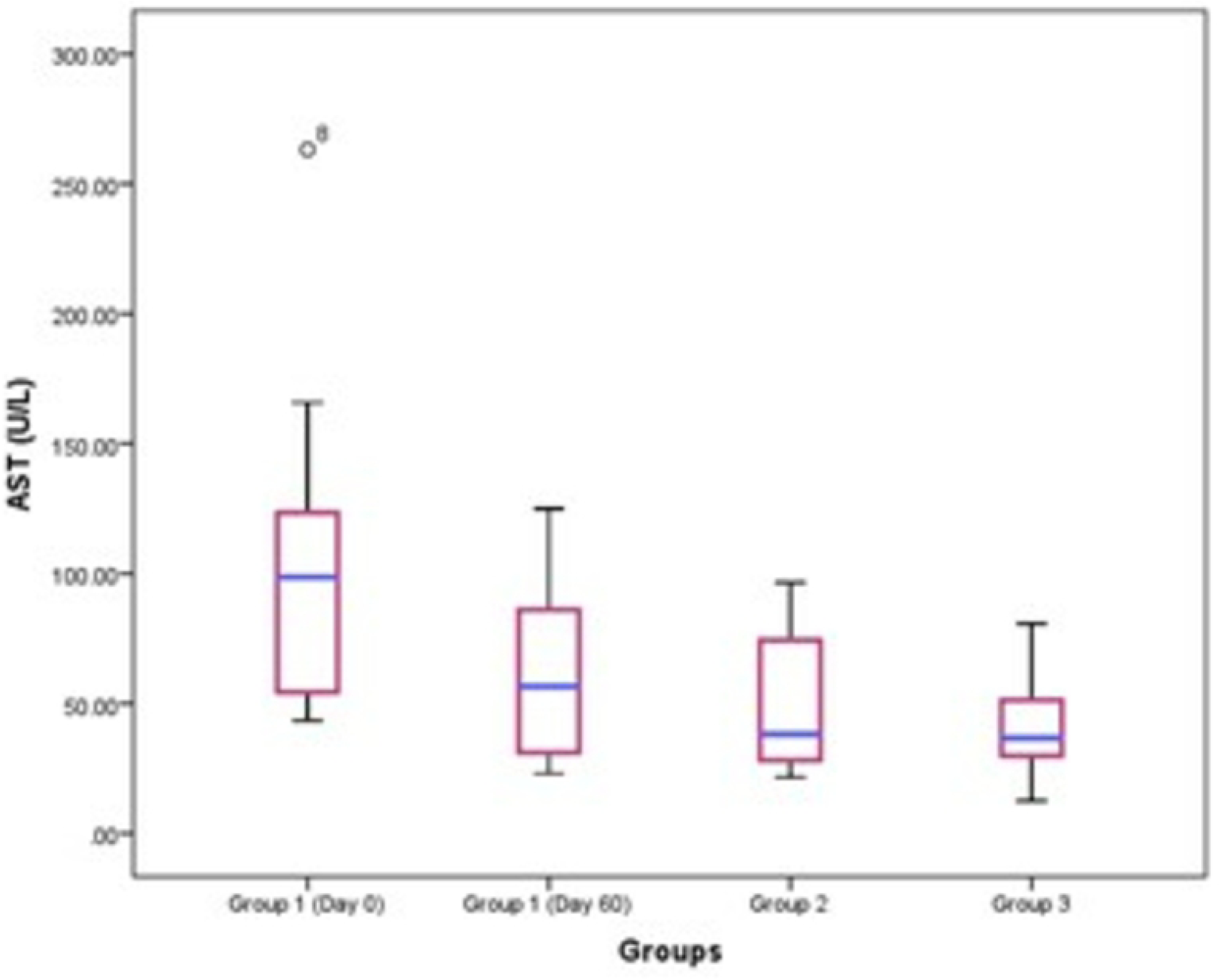
Box plot showing concentration and mean of various serum biochemical parameters in dogs. Concentration of ALP level in dogs. The box plot showing minimum value, the first quartile, the median, the third quartile and the maximum value (vertical number line of Y axis). A value that lies in both extremes of data marked as dots (outlier values).

**Fig 6.**
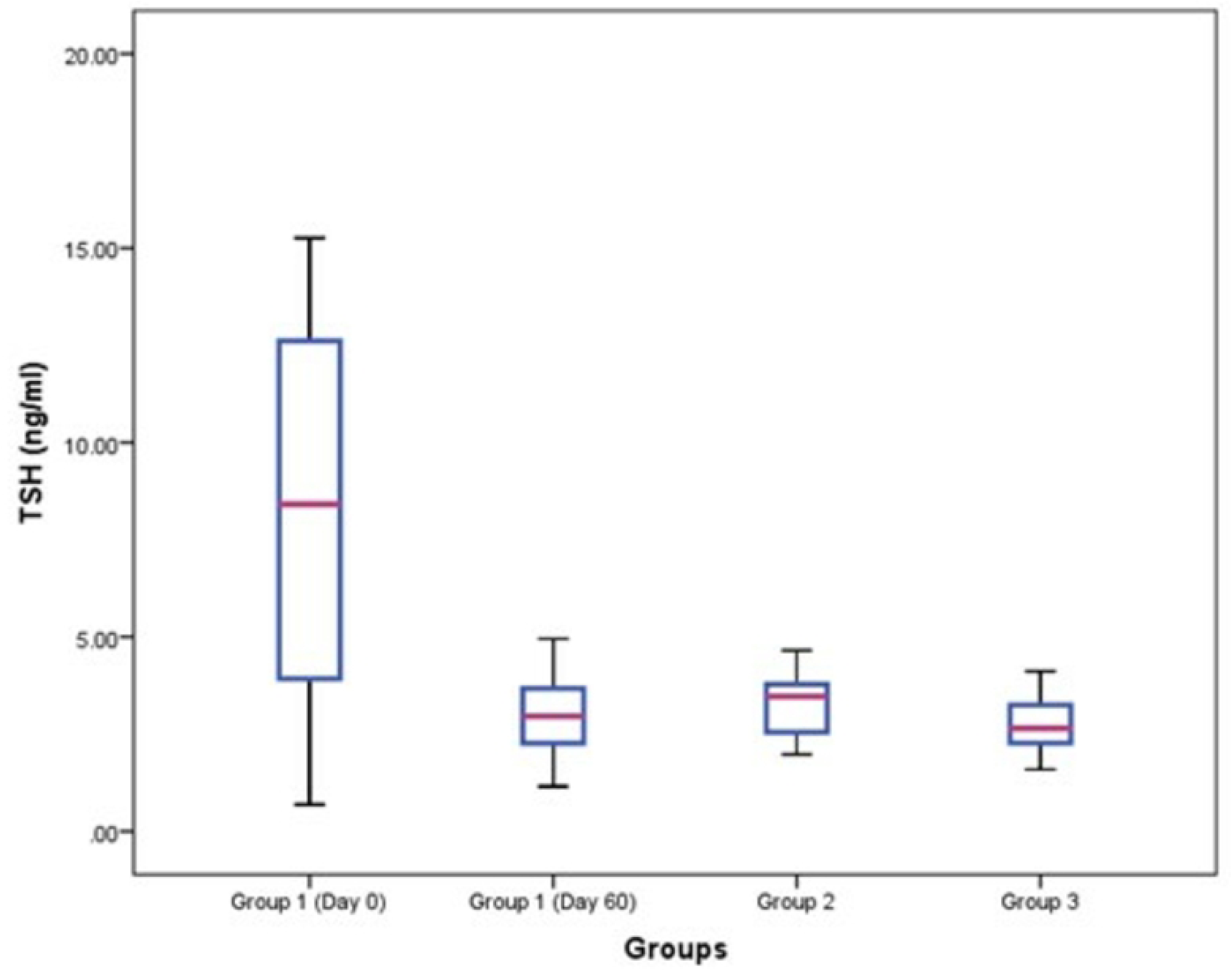
Box plot showing concentration and mean of various serum biochemical parameters in dogs. Concentration of ALT level in dog. The box plot showing minimum value, the first quartile, the median, the third quartile and the maximum value (vertical number line of Y axis). A value that lies in both extremes of data marked as dots (outlier values).

**Fig 7.**
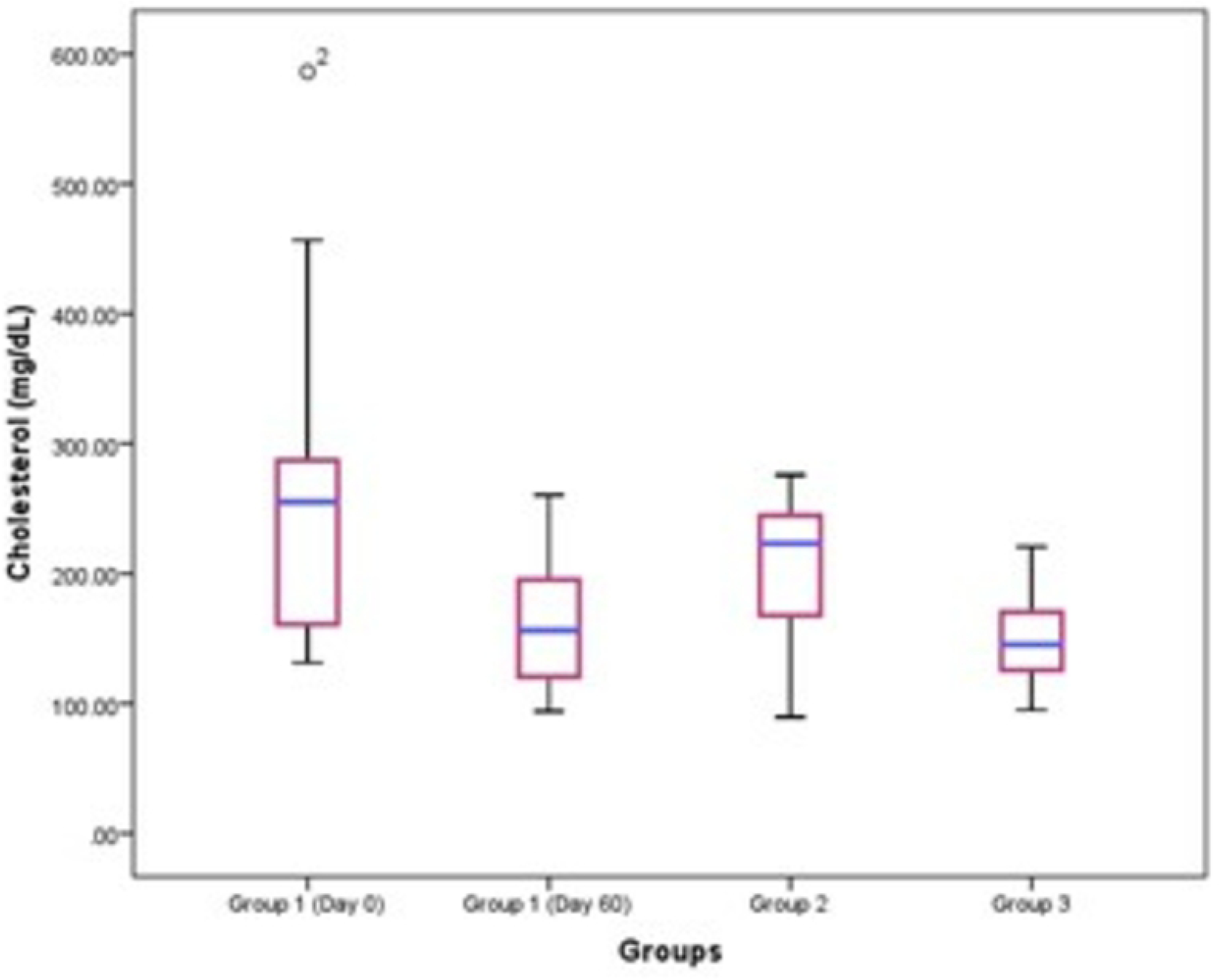
Box plot showing concentration and mean of various serum biochemical parameters in dogs. Concentration of AST level in dogs. The box plot showing minimum value, the first quartile, the median, the third quartile and the maximum value (vertical number line of Y axis). A value that lies in both extremes of data marked as dots (outlier values).

**Fig 8.**
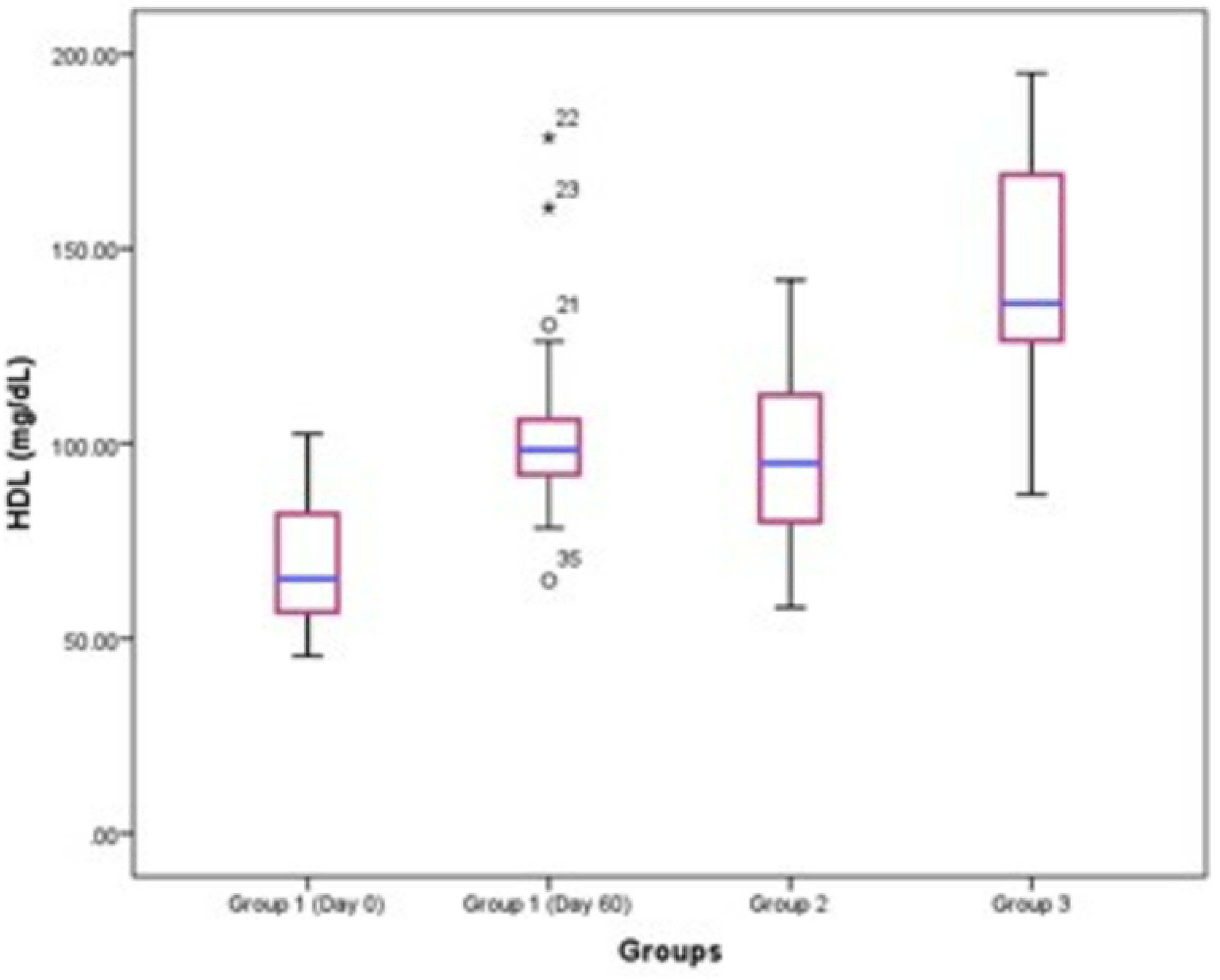
Box plot showing concentration and mean of various serum biochemical parameters in dogs. Concentration of Cholesterol level in dogs. The box plot showing minimum value, the first quartile, the median, the third quartile and the maximum value (vertical number line of Y axis). A value that lies in both extremes of data marked as dots (outlier values).

**Fig 9.**
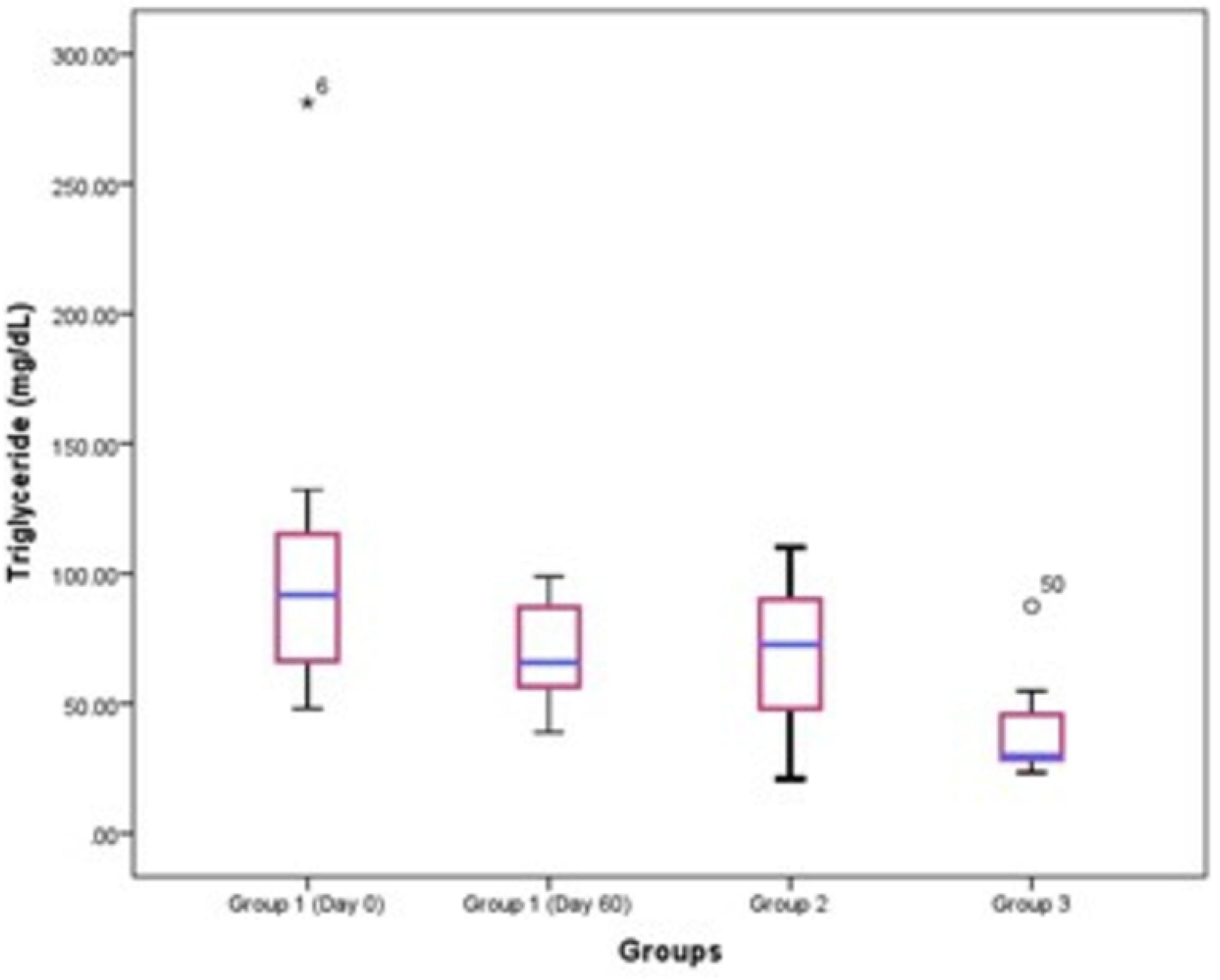
Box plot showing concentration and mean of various serum biochemical parameters in dogs. Concentration of triglyceride level in dogs. The box plot showing minimum value, the first quartile, the median, the third quartile and the maximum value (vertical number line of Y axis). A value that lies in both extremes of data marked as dots (outlier values).

**Fig 10.**
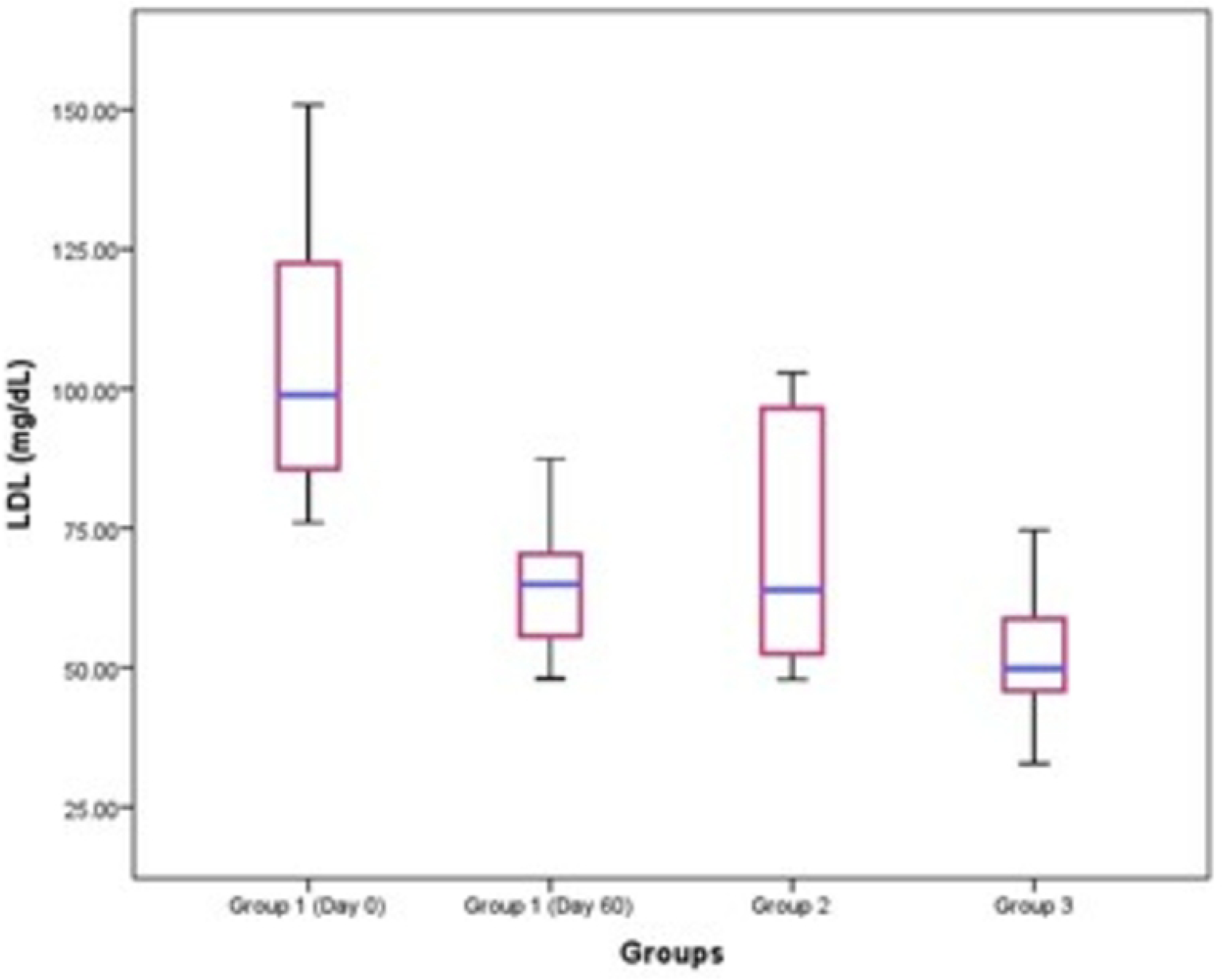
Box plot showing concentration and mean of various serum biochemical parameters in dogs. Concentration of HDL level in dogs. The box plot showing minimum value, the first quartile, the median, the third quartile and the maximum value (vertical number line of Y axis). A value that lies in both extremes of data marked as dots (outlier values).

**Fig 11.**
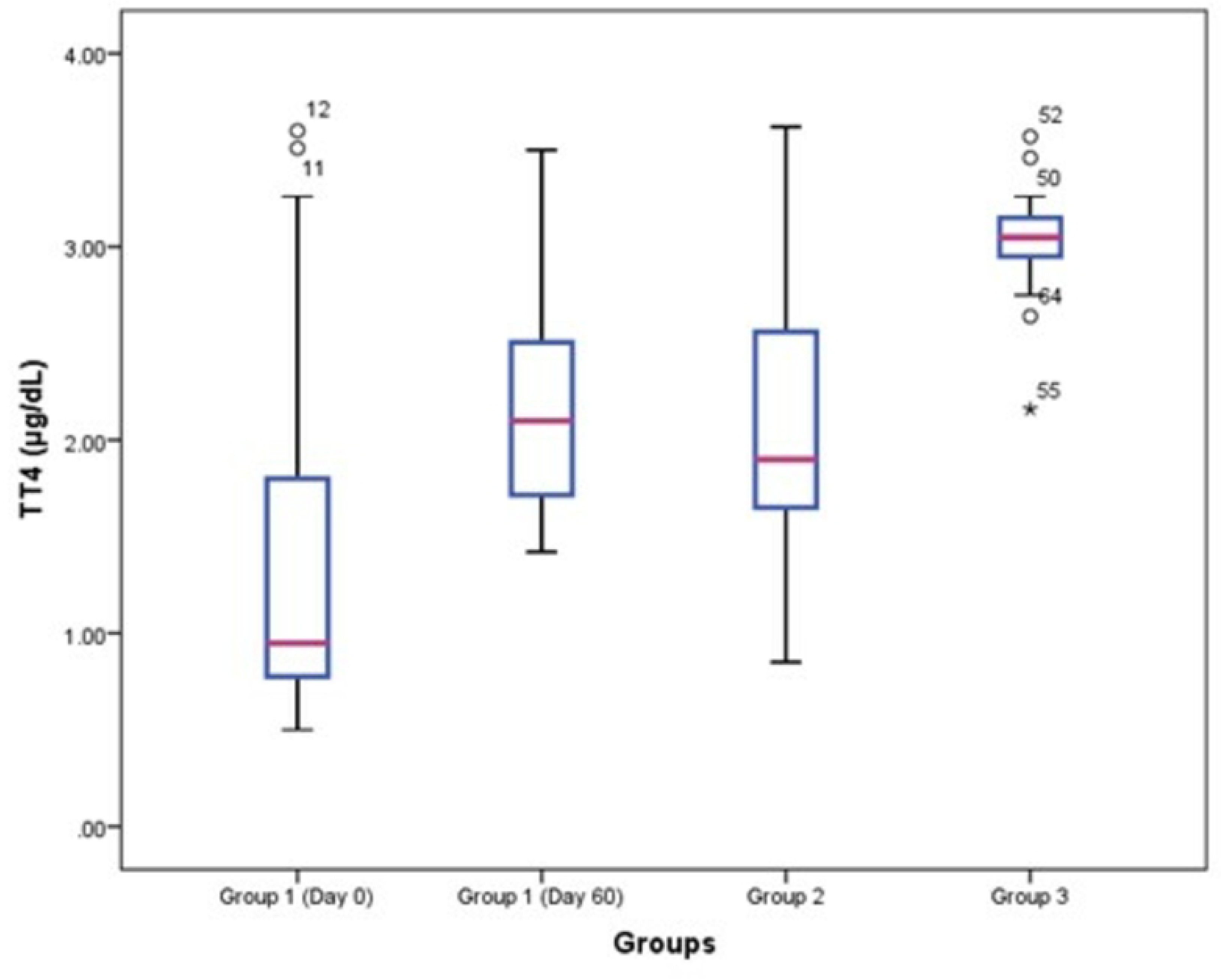
Box plot showing concentration and mean of various serum biochemical parameters in dogs. Concentration of LDL level in dogs. The box plot showing minimum value, the first quartile, the median, the third quartile and the maximum value (vertical number line of Y axis). A value that lies in both extremes of data marked as dots (outlier values).

**Table 3.**
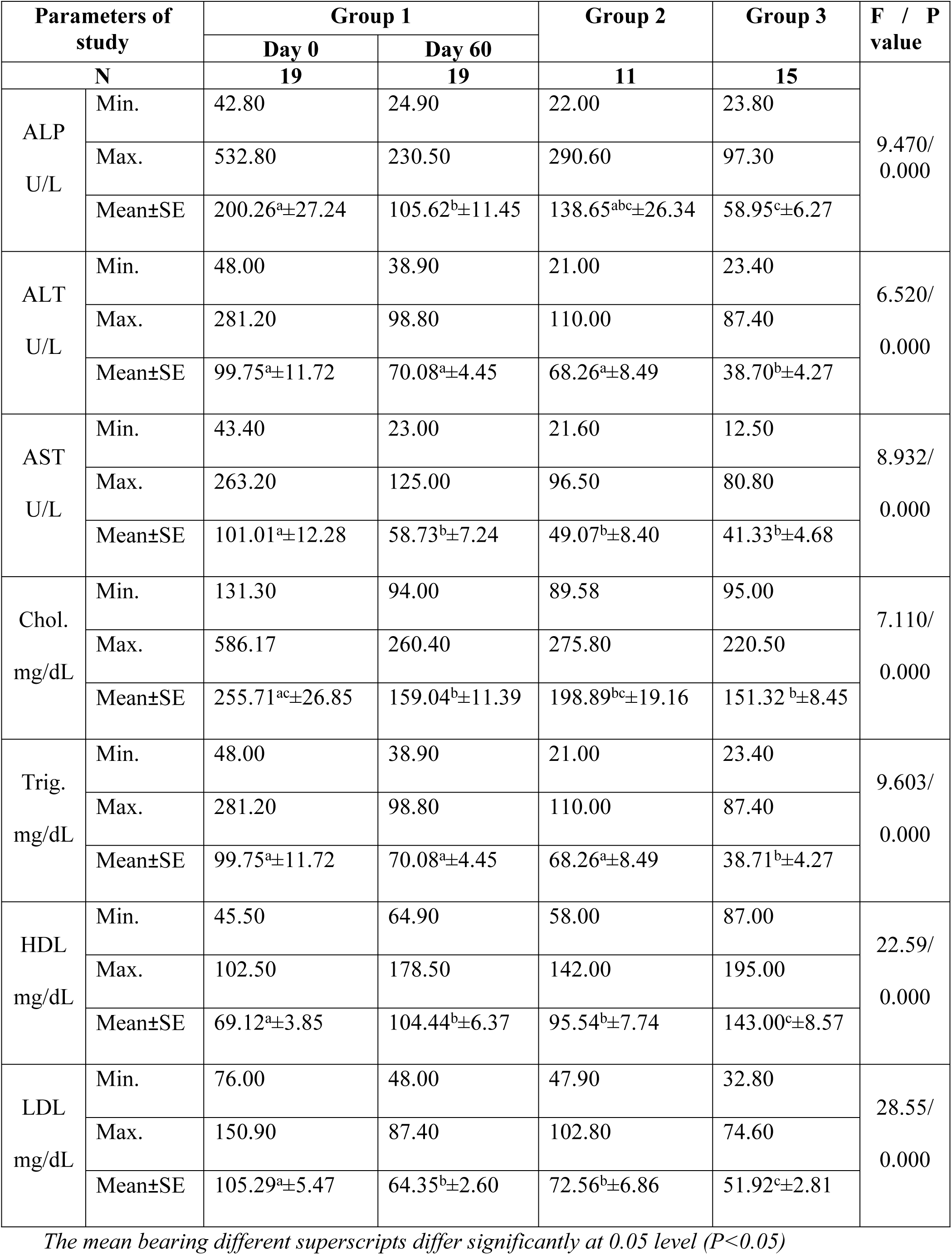
Mean levels of serum biochemical parameters in dogs selected for the study.

#### Assessment of specific antioxidants

Enzymatic antioxidants and peroxidation were measured to rule out oxidative stress in dogs with hypothyroidism, non thyroid illness and euthyroid dogs (Figs 12–22). The activity of catalase, SOD and GSH were significantly lower in group 1 (day 0) compared to other groups, that signified the role of enzymatic antioxidants in hypothyroid dogs. Hypothyroid dogs (group 1) had remarkably higher MDA levels when compared to other groups. Among all oxidative biomarkers lipid peroxidation divulged highly significant variations between groups (P<0.01). Dogs with group 2 showed significant changes in levels of catalase, SOD, GSH and MDA levels with group 1 but not with group 3. Catalase and SOD levels was significantly (P<0.05) differed between hypothyroid dogs on day 0 and day 60, which denoted the importance of therapy in hypothyroid patients. GSH was also lower in hypothyroid dogs on day 0 but whereas reached near normal by day 60. MDA levels was significantly reduced (P<0.05) in hypothyroid dogs after treatment (day 60) (Table 4).

**Fig. 12.**
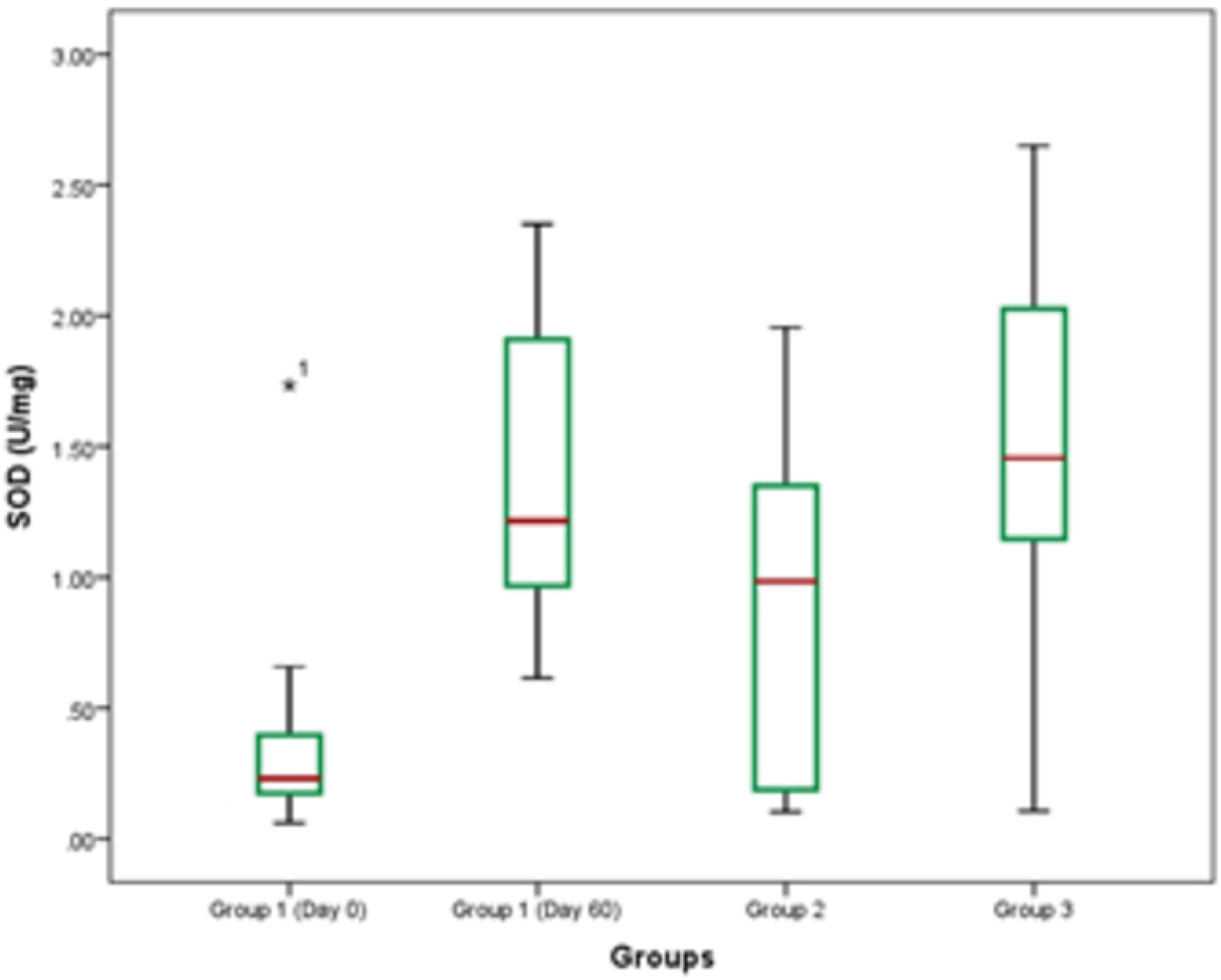
Mean levels of oxidative parameters presented with boxplot in the study population. Concentration Catalase in dogs.

**Fig. 13.**
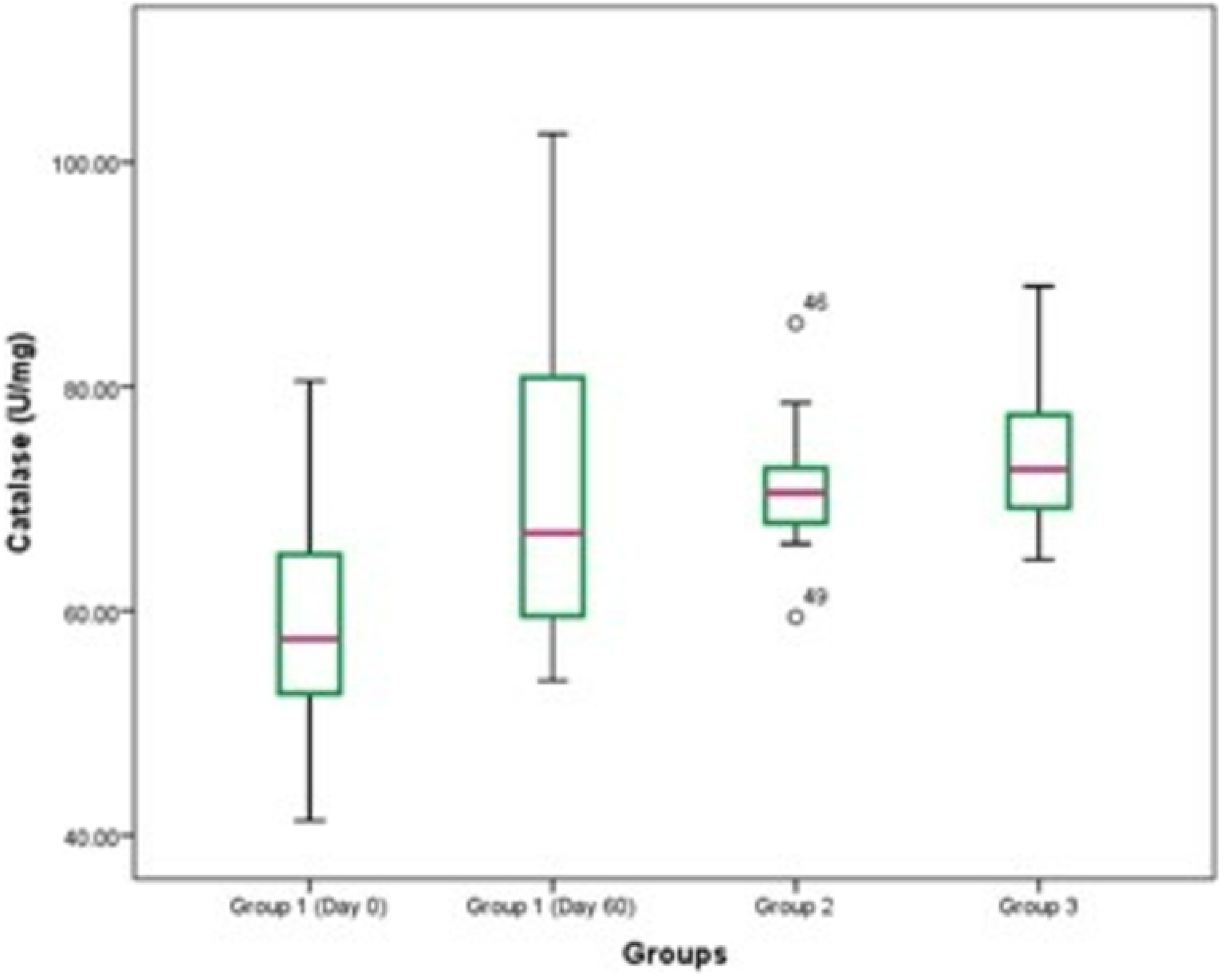
Mean levels of oxidative parameters presented with boxplot in the study population. Concentration of SOD in dogs.

**Fig. 14.**
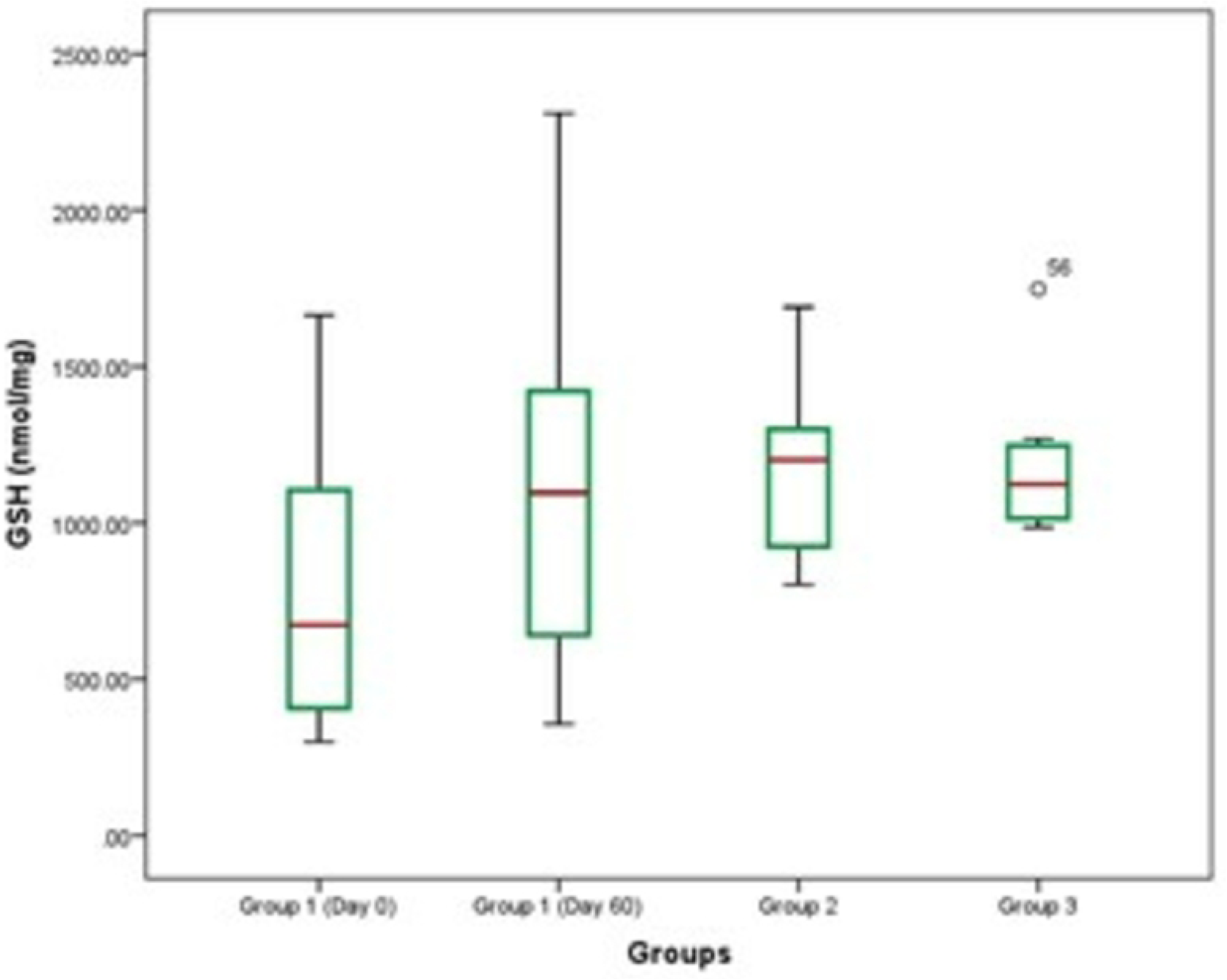
Mean levels of oxidative parameters presented with boxplot in the study population. Concentration of GSH in dogs.

**Fig. 15.**
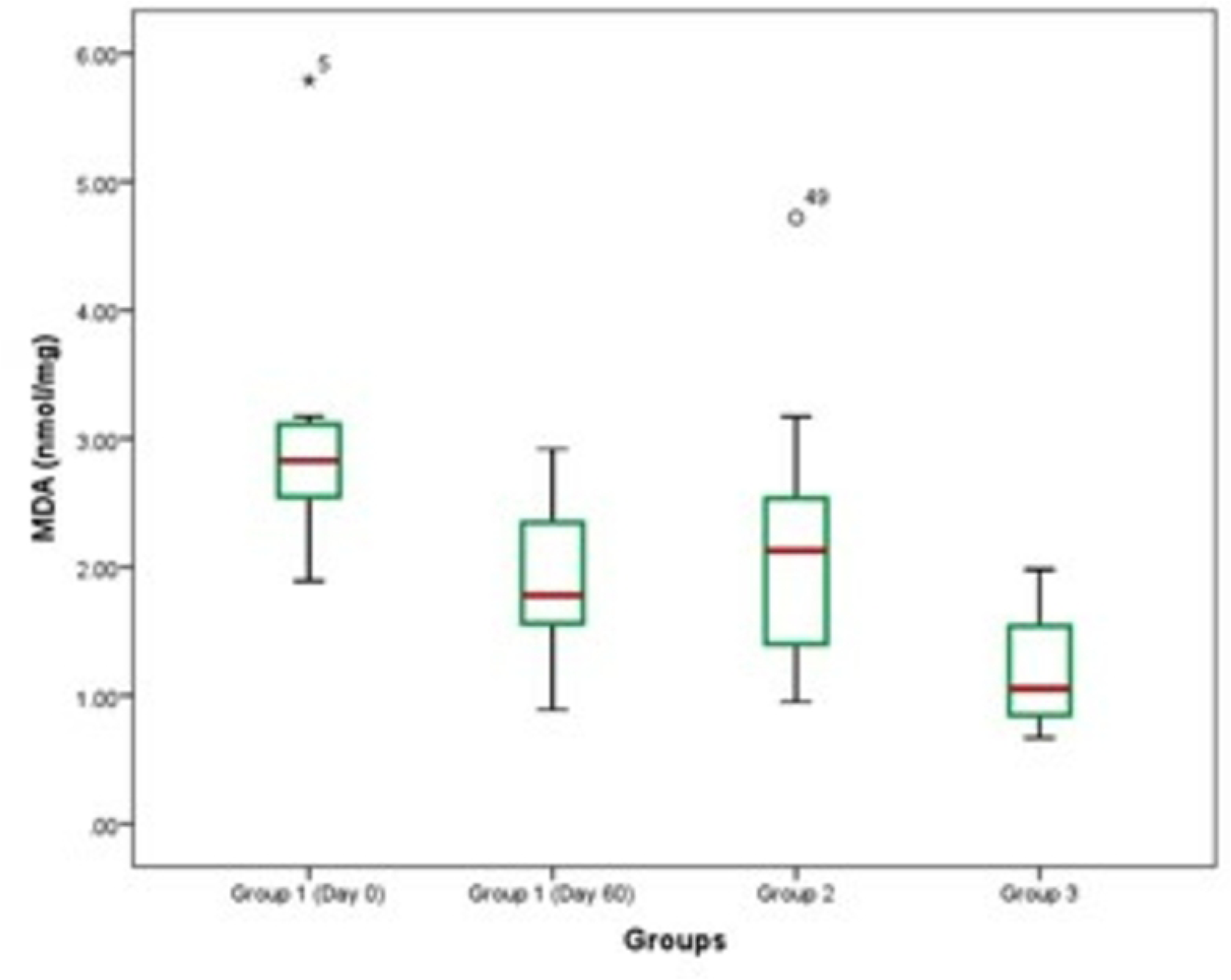
Mean levels of oxidative parameters presented with boxplot in the study population. Concentration of GSH in dogs.

**Table 4.** Mean values of antioxidants in dogs selected for the study.

| Parameters of study |  | Group 1 |  | Group 2 | Group 3 | F value/<br>P value |
| --- | --- | --- | --- | --- | --- | --- |
|  |  | Day 0 | Day 60 |  |  |  |
| N |  | 19 | 19 | 11 | 15 |  |
| Cat.<br>U/mg | Min. | 41.28 | 53.77 | 59.45 | 64.58 | 4.423/<br>0.006 |
|  | Max. | 80.50 | 102.52 | 85.65 | 88.95 |  |
| | Mean±SE | 58.93<sup>a</sup>±2.46 | 70.85<sup>b</sup>±3.36 | 71.23<sup>b</sup>±2.04 | 73.47<sup>b</sup>±1.83 | |
| SOD<br>U/mg | Min. | 0.06 | 0.61 | 0.10 | 0.11 | 17.359<br>/0.000 |
|  | Max. | 1.73 | 2.35 | 1.96 | 2.65 |  |
| | Mean±SE | 0.35<sup>a</sup>±0.08 | 1.38<sup>b</sup>±0.12 | 0.66<sup>c</sup>±0.20 | 1.61<sup>b</sup>±0.18 | |
| GSH<br>nmol/<br>mg | Min. | 298.70 | 356.48 | 801.57 | 985.48 | 2.961/<br>0.039 |
|  | Max. | 1663.64 | 2310.25 | 1690.11 | 1748.00 |  |
| | Mean±SE | 800.79<sup>a</sup>±96.21 | 1113.48<sup>a</sup>±129.13 | 1147.18<sup>b</sup>±81.11 | 1151.16<sup>b</sup>±50.62 | |
| MDA | Min. | 1.89 | 0.89 | 0.95 | 0.67 | 15.533/ |
| nmol/<br>mg | Max. | 5.79 | 2.92 | 4.72 | 1.98 | 0.000 |
|  | Mean±SE | 2.88<sup>a</sup>±0.19 | 1.90<sup>c</sup>±0.14 | 2.20<sup>bc</sup>±0.33 | 1.17<sup>b</sup>±0.11 |  |

The mean bearing different superscripts differs significantly at 0.05 level ( $P < 0.05$ )

#### Correlation study

Correlation analysis was carried out across the obtained results of serum fT4, TT4, cTSH, ALT, AST, ALP, cholesterol, triglycerides, HDL, LDL levels with antioxidant parameters such as catalase, SOD, GSH, MDA levels among hypothyroid, NTI and euthyroid dogs. A highly significant correlation existed between SOD levels and fT4 levels (P<0.01), significant correlation exists between catalase and triglyceride levels (P<0.05) among hypothyroid dogs. Highly significant correlation (P<0.01) exists between levels of fT4, cTSH with MDA among NTI dogs. In euthyroid dogs significant correlation (P<0.05) exists between the levels of cholesterol and TT4 with catalase and MDA (Table 5).

**Table 5.** Correlation coefficients (r) of antioxidant parameters in hypothyroid dogs.

| Paramet. and groups |  | ft4<br>ng/dL<br>(r) | TT4<br>µg/dL<br>(r) | cTSH<br>ng/ml<br>(r) | AST<br>U/L<br>(r) | ALT<br>U/L<br>(r) | ALP<br>U/L<br>(r) | Chol.<br>mg/dL<br>(r) | Trig.<br>mg/dL<br>(r) | HDL<br>mg/dL<br>(r) | LDL<br>mg/dL<br>(r) |
| --- | --- | --- | --- | --- | --- | --- | --- | --- | --- | --- | --- |
| Cat.<br>U/mg | I | -0.245 | 0.067 | 0.324 | -0.207 | -0.218 | 0.003 | 0.126 | -0.465* | -0.245 | -0.285 |
|  | II | 0.100 | 0.395 | 0.296 | -0.502 | 0.015 | 0.562 | 0.435 | 0.053 | 0.057 | 0.087 |
|  | III | -0.331 | -0.561* | 0.127 | 0.237 | 0.513 | 0.313 | -0.533* | -0.156 | 0.011 | -0.334 |
| SOD<br>U/mg | I | 0.879** | -0.095 | -0.166 | -0.023 | -0.142 | -0.229 | 0.010 | -0.126 | 0.266 | -0.227 |
|  | II | -0.076 | -0.411 | -0.483 | 0.266 | 0.052 | -0.325 | -0.193 | 0.340 | 0.282 | 0.431 |
|  | III | -0.042 | -0.576* | -0.222 | 0.323 | 0.138 | -0.364 | 0.074 | -0.036 | -0.501 | 0.154 |
| GSH<br>nmol/<br>mg | I | -0.285 | -0.211 | -0.124 | -0.253 | 0.294 | -0.031 | -0.071 | -0.179 | 0.029 | 0.013 |
|  | II | -0.140 | -0.176 | -0.750 | -0.248 | -0.103 | -0.375 | 0.234 | -0.294 | -0.200 | -0.226 |
|  | III | -0.267 | -0.210 | 0.471 | 0.471 | -0.241 | -0.185 | 0.054 | -0.277 | -0.032 | -0.191 |
| MDA<br>nmol/<br>mg | I | -0.329 | -0.415 | -0.319 | -0.176 | 0.046 | 0.184 | -0.177 | -0.149 | -0.174 | -0.072 |
|  | II | 0.645** | -0.456 | 0.865** | 0.338 | -0.028 | -0.045 | -0.344 | -0.144 | 0.544 | -0.233 |
|  | III | -0.353 | 0.667** | -0.102 | -0.102 | -0.276 | 0.525 | -0.563* | -0.023 | 0.291 | -0.315 |

\*Correlation significant @ 0.05 level, \*\* Correlation significant @ 0.01 level (2 tailed)

### Statistical analysis

IBM SPSS statistics version 21 was used for statistical analysis. One-Way ANOVA, post hoc multiple comparison of unequal variance was done by Games - Howell method (alpha value 0.05). Bi-variate correlation analysis was done by Pearson correlation coefficient method.

## Discussion

The present findings pertaining to hypothyroidism and NTI within the mean age group of 4.9 years (range 1-11 years) and 5.59 years (1-13 years), respectively are in accordance with the findings of previous authors [22, 23]. BCS was higher (6.2) in hypothyroid dogs, when compared to NTI and euthyroid dogs of the present study. Thyroid hormones control body weight, lipid, carbohydrate metabolism, thermogenesis and any alteration in these hormones eventually affect the BCS [24].

Hypothyroid dogs (group 1) had significantly lower fT4 levels (0.56±0.07) and higher mean cTSH levels (8.14±1.17) compared to other groups along with significantly lower level of TT4 compared to group 3. To regulate the levels of thyroxine in hypothyroid dogs, the pituitary gland secretes more TSH. Levels of TSH in serum plays important role in the differentiation of hypothyroidism with NTI and euthyroid dogs. This correlates with the opinion of few authors who stated the importance of cTSH assessment along with TT4 assessment [25, 26] in hypothyroid dogs (TT4 value <12 nmol/L with TSH of >0.5). Whereas, other authors reported fT4 [27] and TT4 [28] assessment as the most sensitive test to rule out hypothyroidism in canines. Present findings are in accordance with the previous authors [29], who reported similar range of serum thyroxine levels, fT4 and TSH levels in hypothyroid and non-thyroidal illness dogs. But the differentiation of non-thyroidal illness from hypothyroidism is still unclear [30].

The significantly elevated levels of total cholesterol and LDL in both hypothyroid and NTI dogs might be associated with obesity, low energy expenditure and increased BCS, that in turn lead to lipid deposition in the liver. Owing to alteration in lipid metabolism associated with hyperlipidaemia in hypothyroid dogs, cholesterol synthesis was increased due to expression of HMD-CoA reductase in the liver [31]. Reduced liver uptake of free fatty acids in hypothyroidism decreased the adipose tissue lipolysis and cholesterol elimination. Further, triglycerides accumulated in the liver and LDL absorption increased due to the persistence of fatty beta oxidation of free fatty acids, that subsequently decreased clearance of triglycerides. Similar findings reported by few authors [32] found increased liver enzymes in hypothyroid dogs as a result of degenerative hepatopathy and myopathy which caused fat infiltration and hyperlipidemia. The authors [33] also documented hypercholesterolemia and hypertriglyceridemia in 75% and 88% of hypothyroid dogs, respectively. Reduced thyroid hormone reduces lipase activity, which mediates fatty acid oxidation for energy production. Observational studies in human patients proved that 30% of patients with overt hypothyroidism had dyslipidemia and triglyceride accumulation in the liver, which is a risk factor for rising levels of LDL and total cholesterol [34]. Previous studies reported [35] the positive reversal of hyperlipidemia in hypothyroid patients with exception of underlying hyperlipidemia.

According to the authors knowledge there were few studies on oxidative parameters in hypothyroid dogs and hence the current investigation reports put on record for the first time in India about oxidative stress in hypothyroid dogs, dogs with NTI and euthyroid dogs. Different biomarkers were proposed by various authors, MDA is one such indicator of lipid peroxidation [36,37]. In addition to that, enzymatic biomarkers including SOD, catalase, and GSH were evaluated in the present investigation to measure oxidative stress. The changed equilibrium between pro and antioxidants were identified in the current investigation by measuring alterations in enzymatic antioxidants and lipid peroxidation in hypothyroid dogs. When pro-oxidants and antioxidants are in an unbalanced ratio, antioxidant defence failed and lead to oxidative stress.

Significant finding of the current investigation was high MDA levels in hypothyroidism and NTI dogs when compared to the euthyroid dogs. The mean level of MDA was greater in the hypothyroid dogs (2.88±0.19) (P<0.05), which revealed severe oxidative stress in hypothyroid dogs. The findings of which were similar to the earlier authors [36–38] and whereas, others [39] documented decreased lipid peroxidation in hypothyroid rats. According to the current study, levothyroxine supplementation might have helped to enhance oxidative status in dogs with hypothyroidism. It markedly lowered the MDA levels and increased the catalase, SOD and GSH activity. Similar findings have been reported earlier [7, 40–42] suggesting the increased oxidative stress in canine hypothyroidism. In human subjects the oxidative status and levothyroxine correlation was studied and reported [8]. However, some previous studies also showed inconsistent results [6,43].

The current study further reported a reduced catalase levels in hypothyroid dogs compared to NTI and euthyroid dogs, that could be due to reduced antioxidant defence in hypothyroid dogs, which was restored by levothyroxine therapy. Some human studies were contradictory as well. However, some studies in human primary hypothyroid patients reported higher catalase activity [6,8] but some have reported lower levels [44].

The activity of SOD was significantly lower in hypothyroid dogs when compared to NTI and euthyroid dogs that was significantly elevated to normal with levothyroxine supplementation. Similar reports were also published [37,44] in human hypothyroid patients. Other research studies found no appreciable difference in SOD activity between hypothyroidism and euthyroidism [6,8] which may be due to variations in the stage of hypothyroidism and concurrent illness. Our study identified significant difference of SOD activity levels between hypothyroidism dogs prior to and during levothyroxine therapy, which signifies the SOD activity in antioxidant defence.

## Conclusion

The current investigation established the fact that the dogs with hypothyroidism showed the evidence of oxidative stress. This study mainly demonstrated the changes in antioxidant parameters such as catalase, SOD, GSH and MDA in hypothyroid dogs compared to NTI and euthyroid dogs. Hypercholesterolemia, hypertriglyceridemia, elevated levels of LDL, ALP, AST, ALT and lower level of HDL were documented in hypothyroid dogs. All the biomarkers have shown significant alteration in dogs with hypothyroidism and levothyroxine therapy restored the oxidation state. So, these biomarkers can be used as a useful measure to monitor oxidative stress in hypothyroid dogs and to evaluate the therapeutic efficacy. These findings can be a torch bearer for future research in canine subject.

